# Dynamic evolution of recently duplicated genes in *Caenorhabditis elegans*

**DOI:** 10.1101/2022.03.10.483751

**Authors:** Fuqiang Ma, Chun Yin Lau, Chaogu Zheng

## Abstract

As a major origin of evolutionary novelties, gene duplication is a widespread phenomenon across species. However, the evolutionary force that determines the fate of duplicate genes is still under debate. Here, we studied the functional evolution of duplicate genes at both macroevolution and microevolution scales using the genomic sequences of eleven *Caenorhabditis* species and 773 *C. elegans* wild isolates. We found that compared to older duplicate genes and single-copy genes, recently duplicated gene copies showed rapid turnover, large genetic diversity, and signs of balancing and positive selection within the species. Young duplicate genes have low basal expression restricted to a few tissues but show highly responsive expression towards pathogenic infections. Recently duplicated genes are enriched in chemosensory perception, protein degradation, and innate immunity, implicating their functions in enhancing adaptability to external perturbations. Importantly, we found that young duplicate genes are rarely essential, while old duplicate genes have the same level of essentiality as singletons, suggesting that essentiality develops over a long time. Together, our work in *C. elegans* demonstrates that natural selection shapes the dynamic evolutionary trajectory of duplicate genes.

**Significance:** The “evolution by gene duplication” theory suggests that gene duplications provide the genetic materials for mutation and selection to act upon, expand the repertoire of molecular functions, and enable evolutionary novelty. Although various models were proposed to describe the fate of duplicate genes, empirical evidence for these models is limited. We analyzed gene duplications in eleven nematode *Caenorhabditis* species and studied the intraspecific variation of these duplicate genes among *C. elegans* wild strains. We found that compared to older duplicate gens and single-copy genes, recently duplicated genes show rapid turnover, large genetic diversity, and strong signs of balancing and positive selection but rarely develop essential functions. Our results describe the evolutionary trajectory of duplicate genes shaped by natural selection.

## Introduction

Gene duplication is a major mechanism to produce new genetic materials, and the duplicate genes can, in theory, undergo subfunctionalization, neofunctionalization, and degeneration (1-4). The evolutionary mechanism that drives the preservation of duplicate genes is under intense research. Although the classical duplication-degeneration-complementation (DDC) model predicts that duplicate genes are preserved by the partitioning of ancestral functions (2), more recent studies found that positive selection may have driven the retention of duplicate genes in Drosophila (5) and mammals (6) mostly through neofunctionalization. Intriguingly, Chen et al. (7) found that in Drosophila 30% of the young genes, arising from recent duplication events in the last 35 million years, integrated into vital pathways and acquired essential functions, suggesting that purifying selection may quickly become the driving force to preserve new duplicate genes. More empirical evidence will be needed to understand whether and how natural selection shapes the evolutionary trajectory of duplicate genes. Moreover, previous studies addressing this question mostly compared the sequences of paralogs or the sequences of homologs between species; however, we reason that examining the intraspecific variation and selection of the recently duplicated genes at the population level may provide fresh insight into the evolutionary fate of duplicate genes.

Another important question that remains unclear is whether similar selection force operates on both young and old duplicate genes. One compelling hypothesis is that newly duplicated genes are first retained by positive selection, then gradually evolve essential functions, and are eventually preserved by purifying selection (8, 9). However, information from genome-wide comparison of genes duplicated at different time points in evolution is quite limited. Furthermore, the function of the young duplicate genes and the biological pathways they are involved in are largely unknown.

In this study, we address the above questions by analyzing the functional evolution of duplicate genes in the nematode *Caenorhabditis elegans* at both the macroevolution (between species) and microevolution (within species) scales. *C. elegans*, a widely used model organism for biological research, has emerged as an important species for evolutionary studies, thanks to the availability of the genomic sequences of >700 wild strains (10) and dozens of related nematode species (11, 12). Importantly, *C. elegans* has a high spontaneous gene duplication rate (13), making it a good species to study the fate of duplicate genes. Although pioneering works have identified duplications in specific gene families in *C. elegans* (14, 15), analyzed the structural heterogeneity between duplicate genes (16), and studied the functional redundancy of paralogs (17, 18), a genome-wide analysis of the evolutionary fate of duplicate genes is still missing.

In this study, we first identified *C. elegans* genes arising from recent and old duplications by comparing the genomes of eleven *Caenorhabditis* species and then analyzed the intraspecific variation of these duplicate genes among 773 *C. elegans* wild isolates. We found that the young duplicate genes undergo dynamic interspecific and intraspecific evolution—showing much larger nucleotide polymorphisms and copy number variations than single-copy genes (termed singletons) and old duplicate genes, signs of balancing and positive selection, and fixation in certain genetic groups. Genes involved in chemosensory perception, protein degradation, and innate immunity were enriched in the recently duplicated genes, suggesting possible involvement of these pathways in adaptation. The expression of recently duplicated genes was restricted to a few tissues and showed high plasticity in response to stress conditions and pathogens; the expression patterns of paralogs also diverged significantly at the single cell level. Importantly, we found that the old duplicate genes and singletons share similar levels of essentiality and are under purifying selection, whereas young duplicate genes rarely become essential, suggesting that it takes a long time to develop essential functions. Overall, our systematic analysis in *C. elegans* provides an example to support that natural selection plays a vital role in shaping the evolution of duplicate genes.

## Results

### Gene duplication shapes genomic evolution in eleven *Caenorhabditis* species

To analyze gene duplication in the evolution of genomes in the *Caenorhabditis* genus, we collected the protein sequences from eleven *Caenorhabditis* species that have high-quality genomic data (Table S1; See Methods for details) from WormBase ParaSite and constructed orthogroups (OGs) among the homologous genes using Orthofinder2 (19). Most (∼90% or above) of the genes were included in OGs for each species (Fig. 1A; Table S2). By comparing the gene tree of each OG and the species tree under a duplication-loss-coalescence (DLC) model (20), we identified duplication events and genes derived from duplications at each branch of the phylogenetic tree (Table S3). We then grouped the duplicated genes by their origin from recent (within 60 million years or mys), late (60-130 mys ago), or ancient (>130 mys ago) duplications; the divergence times were calculated based on the information on TimeTree (timetree.org; see Methods). Considerably more genes were duplicated in recent times than in the late and ancient times for all species, ranging from 14% to 34% of the genomes (Fig. 1A).

**Figure 1.**
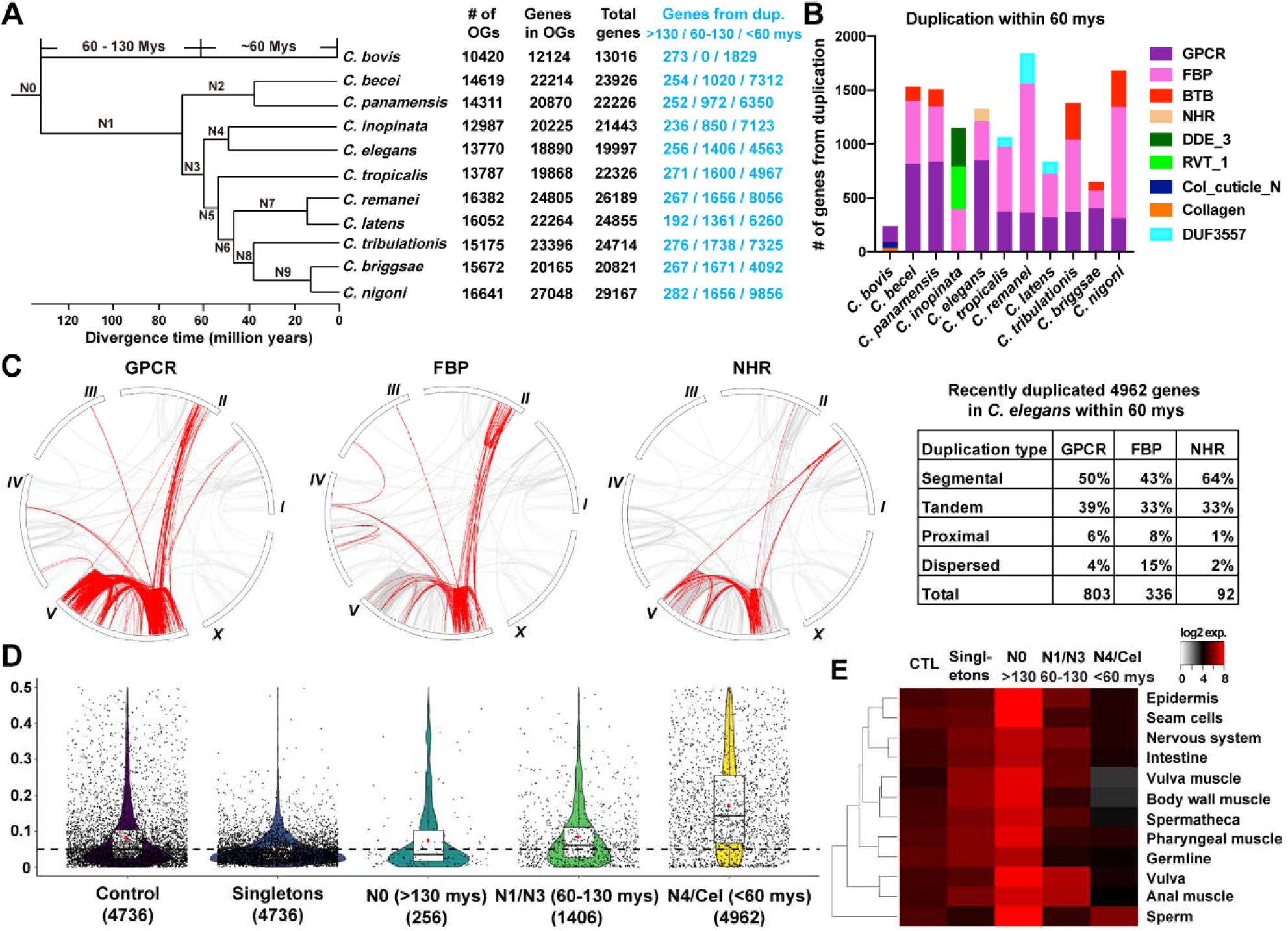
Duplication characteristics among eleven *Caenorhabditis* species. (A) The number of genes in the orthogroups (OGs) and genes duplicated at different times for each *Caenorhabditis* species identified by Duplication Loss Coalescence model. (B) The top three domains for the genes duplicated within 60 million years for each species. (C) Circle plots showing the collinearity of recently duplicated GPCR, FBP and NHR genes (red lines) over the grey collinearity background in *C. elegans*. The duplication types were assigned according to the output of MCScanX. (D) The violin plot showed the evolutionary rate (dN/dS) for *C. elegans* singletons and duplicate genes at N0, N1/N3, and N4/Cel branches. The control showed the results of 4506 randomly chosen genes from the genome. Genes with dN or dS value smaller than 0.0005 or equal to 999 were removed. Horizontal dashed line indicates dN/dS = 0.05. (E) Heatmap for mean tissue expression of singletons and genes duplicated at N0, N1/N3 and N4/Cel (expression data were obtained from CeNGEN).

Using PfamScan (21), we searched for domain information of the recently duplicated genes (within 60 mys) and found that several domains were common among these genes, including G protein-coupled receptors (GPCRs), F-box domain (FBPs), BTB domain, DUF3557 domain, etc., suggesting that similar families of genes were duplicated in different *Caenorhabditis* species (Fig. 1B). Interestingly, *C. inopinata* had significant duplications in genes involved in DNA transposition (DDE_3 domain) and retrotransposition (RVT_1 domain), which did not occur in the sister species, *C. elegans*.

Focusing on *C. elegans*, we generated a list of 4,962 recently duplicated genes (labeled as N4/Cel in Table S4) that include the 67 genes duplicated at the N4 branch, 4,496 genes duplicated at the *C. elegans* terminal branch (Fig. 1A), and 399 genes found in *C. elegans*-specific OGs that only contained 2 or 3 genes and were thus excluded from the DLC model. We also identified 256 and 1,406 genes originated from ancient (at N0 branch) and late (at N1 and N3 branches) duplications, respectively. As controls, we identified 4,736 singletons from the OGs that had only one homolog from each of the eleven species (Table S4).

To visualize the evolutionary history of the recently duplicated genes, we made collinearity plots for the top three duplicate gene families, including the GPCRs, FBPs, and nuclear hormone receptors (NHRs) containing the zf-C4 domain (Fig. 1C). In all three cases, most duplications were segmental or tandem, and the duplication of genomic blocks within the chromosome V was a major driving force for gene duplications. In addition, we observed duplications between the left arm of chromosome II and the right arm of chromosome V for FBPs, between the right arm of chromosome I and the chromosome V for NHRs, and both for GPCRs (Fig. 1C).

To understand the sequence divergence and the evolutionary constraints of the duplicate genes, we calculated the dN/dS ratio (the number of nonsynonymous substitutions per nonsynonymous site over the number of synonymous substitutions per synonymous site) for homologous genes in the same OGs and found that recently duplicated genes had excessive nonsynonymous variations compared to singletons (Fig. 1D). Interestingly, genes originating from older duplications (N1/N3) had lower dN/dS ratios than the recently duplicated genes (N4/Cel), suggesting stronger evolutionary constraints. Thus, young duplicate genes may be functionally more divergent than singletons and old duplicate genes and may be positively selected.

We examined the expression levels of duplicate genes in different tissues using the recent single-cell transcriptomic data (22). Genes derived from recent (N4/Cel) duplications showed expression in fewer tissues and had weaker expression in most tissues than singletons and genes derived from older duplications (at the N0 and N1/N3 branches) (Fig. 1E). This result is consistent with weaker purifying selection (bigger dN/dS) in recently duplicated genes and matches previous observations in other organisms (23). Strikingly, *C. elegans* genes arising from recent duplication showed highly enriched expression in sperms (Fig. 1E), which supports the “out of the testis” hypothesis that selective pressures in the male germline drives the creation and fixation of new genes (24). Our results and recent work in another nematode *Pristionchus pacificus* (25) extended this idea beyond Drosophila and mammals to androdioecious species.

### Expansion of specific gene families through duplications

We next identified the gene families that underwent significant expansion through duplication by applying the Computational Analysis of gene Family Evolution (CAFE) algorithm, which uses a stochastic birth and death process to model the change of gene family sizes over the phylogeny of the species (26). This analysis uncovered 922 and 299 OGs that were significantly expanded and contracted, respectively, at the species level (Fig. 2A). Among the eleven species, *C. nigoni* had the largest number (268) of expanded OGs, whereas its sister species *C. briggsae* had the largest number (122) of contracted OGs, which explains the differences in their total gene numbers (Fig. 1A).

**Figure 2.**
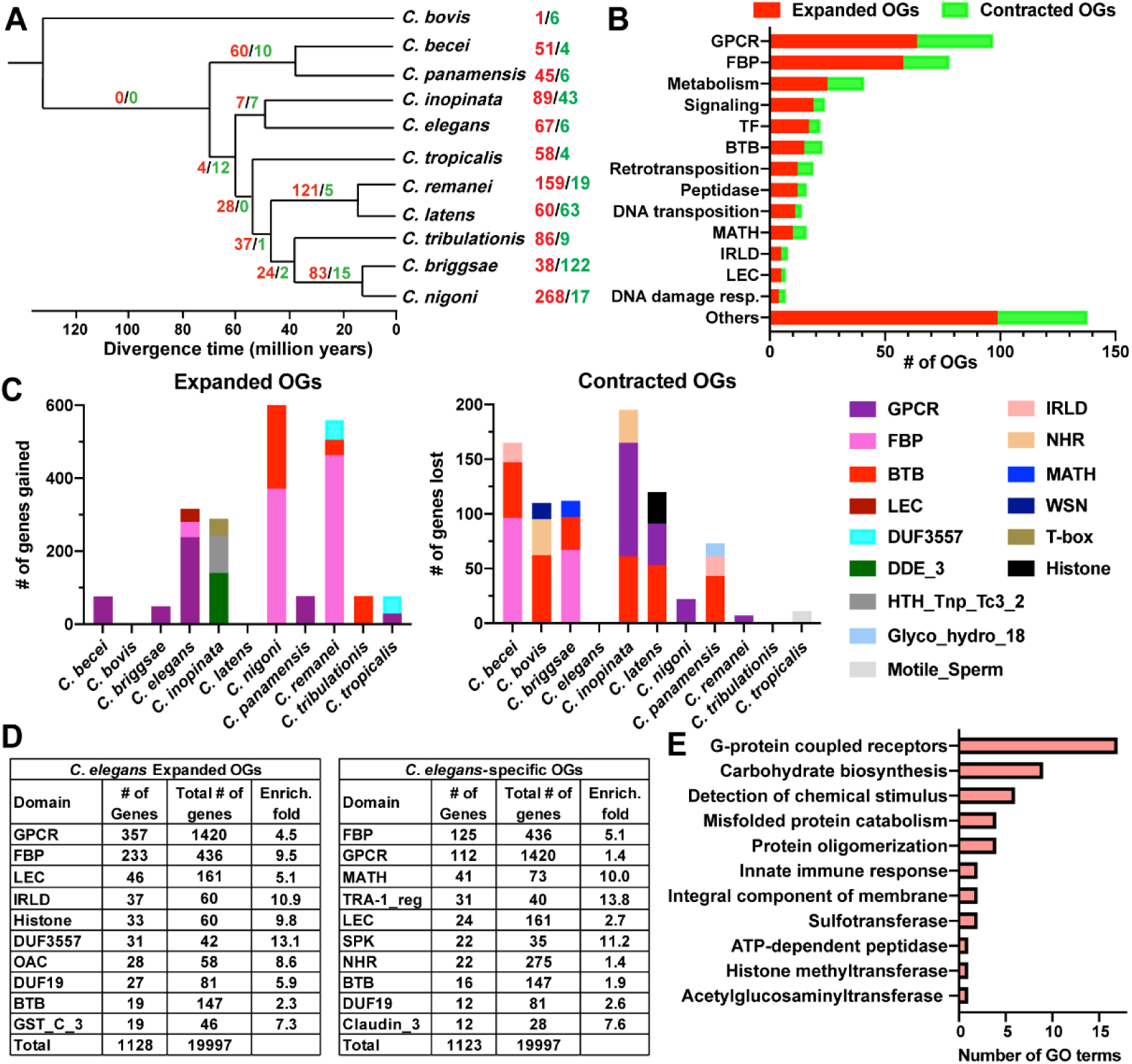
Expansion of specific gene families through duplications. (A) The number of significantly expanded (red) and contracted (green) OGs in the eleven *Caenorhabditis* species and inner branches. (B) The number of expanded (red) and contracted (green) OGs with specific domains or functions in the eleven species. The grouping of OGs was shown in Table S5. (C) The number of genes gained and lost in the significantly expanded and contracted OGs within 60 million years, respectively, were shown for the top three affected gene families. The number of genes gained and lost were calculated as the difference between the number of genes in one species and the average number of genes among the other ten species. Gained and lost domains with less than thirty and ten genes, respectively, were not shown. (D) The number and enrichment fold of genes with specific domains (top ten domains) in *C. elegans* expanded and specific OGs. The total number of genes with specific domain were found by searching all *C. elegans* proteins with PfamScan. (E) The enrichment modules for GO functional category for the 2,256 *C. elegans* expanded and specific genes.

We then used PfamScan to search for domain information and found that the significantly expanded or contracted OGs share certain molecular structure or biological functions (Fig. 2B; Table S5). For example, the two biggest gene families that were under expansion were GPCRs (64 OGs) and FBPs (58 OGs). Proteins with MATH and BTB domains were also commonly found in the expanded OGs. GPCRs detect extracellular cues and transduce signals across the membrane; both FBP and MATH-BTB proteins act as substrate-specific adaptors for E3 ubiquitin ligase and function in protein degradation. In addition, genes involved in DNA transposition, retrotransposition, metabolism, and signaling, as well as TFs, helicases, and peptidases are found in OGs with species-level expansion (Fig. 2B; Table S5). Interestingly, similar domains were also found in the contracted OGs, suggesting that these gene families have undergone dynamic evolution in *Caenorhabditis*, expanding in one species while contracting in another. Different *Caenorhabditis* species had expanded and contracted OGs with distinct domains (Fig. 2C). For example, *C. nigoni* and *C. remanei* significantly expanded the FBP gene families, while the most expanded OGs in *C. becei, C. briggsae, C. elegans*, and *C. panamensis* contained GPCRs.

Next, we focused on the 1,128 genes in the 71 *C. elegans* expanded OGs and the 1,123 genes in 264 *C. elegans*-specific OGs, which were excluded from the above CAFE analysis because they only contained genes from one species (Table S6). 95% of the 2,251 genes in the *C. elegans* expanded and specific OGs were found in the 4,962 recently duplicated genes, suggesting that two independent methods identified the same duplicate genes. Domain analysis showed that the GPCR, FBP, LEC (containing the Lectin_C domain), IRLD (containing the Recep_L_domain), histones, DUF3557, OAC (containing the Acyl_transf_3 domain), and other gene families were highly enriched in *C. elegans* expanded OGs. GPCR, FBP, MATH, LEC, NHR, and genes containing the TRA-1_regulated domains were enriched in *C. elegans*-specific OGs (Fig. 2D; Table S6). Gene ontology (GO) analysis found the enrichment of these genes in functional GO terms consistent with domain characterization (e.g., OAC for carbohydrate biosynthesis, GPCR for detection of chemical stimulus, LEC and FBP for innate immune response, etc.). The expansion of specific gene families involved in sensory perception, innate immunity, and other pathways might contribute to adaptation during the evolution of *C. elegans*.

### Positive selection and functions of duplicate genes in *C. elegans* expanded and specific OGs

*C. elegans*-specific OGs contained only *C. elegans* genes, while the *C. elegans* expanded OGs showed varying degrees of expansion in *C. elegans* for different gene families (Fig. 3A and Fig. S1). For instance, the four representative FBP OGs are made of more than 95% *C. elegans* genes, whereas the OAC OG contains less than 20% *C. elegans* genes.

**Figure 3.**
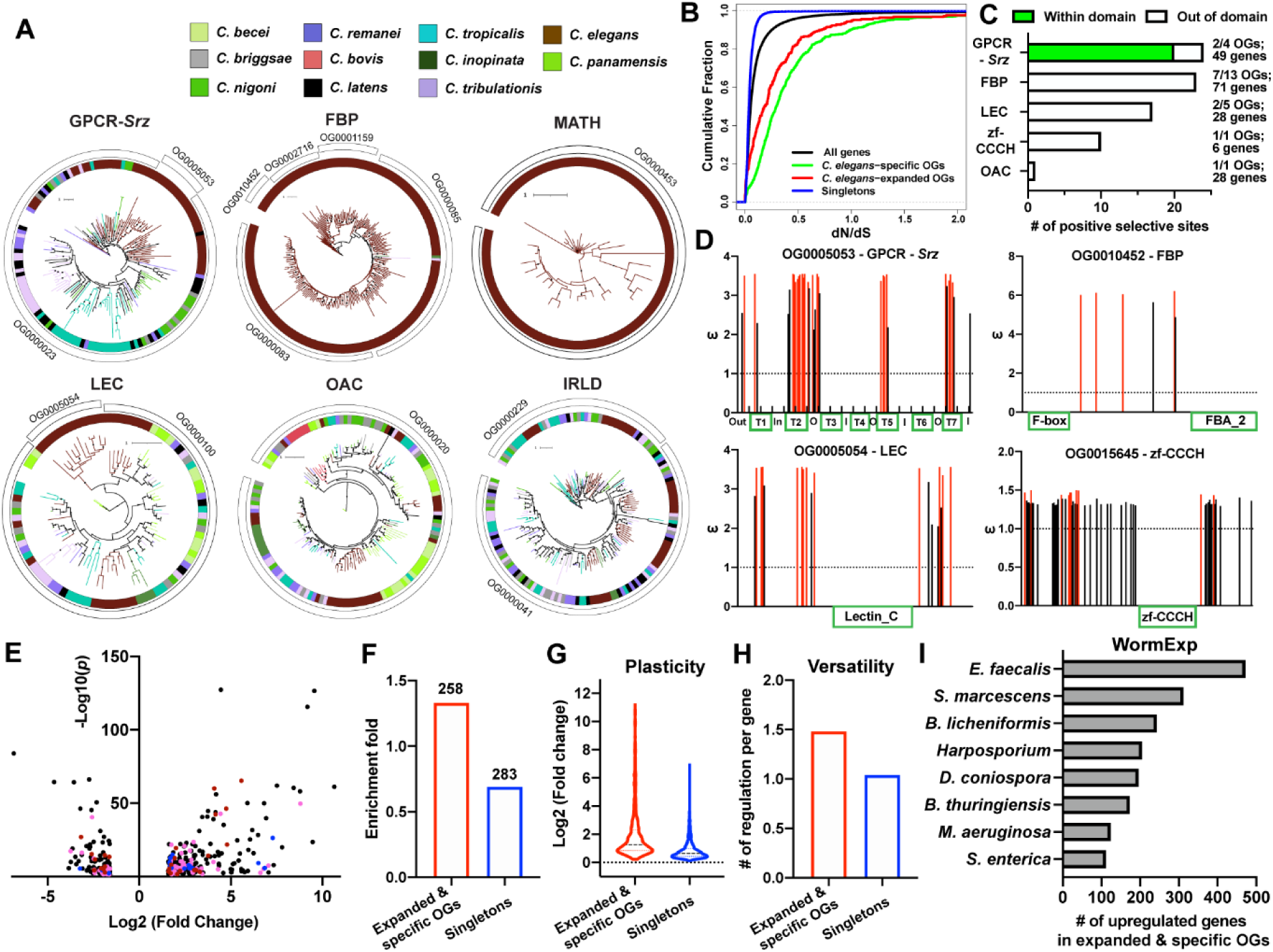
Positive selection and responsive expression on *C. elegans* expanded and specific genes. (A) The phylogenetic trees constructed using maximum likelihood method with one thousand bootstraps for the *C. elegans* expanded or specific OGs with specific domains. OGs sharing the same domain were merged for tree construction. Branches with more than 80% bootstrap support were marked by black point (zoom in to see). (B) The cumulative distribution of evolutionary rate (dN/dS) for genes in *C. elegans* specific and expanded OGs, singletons, and all genes. (C) The number of positively selected sites and the number of OGs with the sites in gene families from the *C. elegans* specific and expanded OGs. The sites were identified with Bayesian posterior probability of positive selection > 0.9. For GPCRs, the domain means the transmembrane helices. (D) The distribution of positively selected sites (dN/dS > 1) along the codons of four representative OGs in *C. elegans*. Vertical red lines indicate sites with ω values with posterior probability over 0.9. The green box indicates domain position. The horizontal dash line indicates dN/dS = 1. (E) The volcano plot showing the differential expression (fold change > 3 or < 1/3) of duplicate genes in the *C. elegans* expanded and specific OGs upon exposure to heat shock and five pathogens. Pink, FBP genes; blue, MATH genes; red, LEC genes. (F) The number (on top of the bars) and enrichment fold of the duplicate genes and singletons among the significantly (fold change >3) upregulated genes. (G) The dynamic range of fold changes for these genes among the upregulated genes (fold change > 0). (H) The average number of stress conditions that induced significant upregulation (fold change > 3) of the genes. (I) The number of genes in *C. elegans* expanded and specific OGs were enriched in genes upregulated by infection of several pathogens based on the analysis of WormExp under the “Microbe” category.

Genes from the *C. elegans* expanded and specific OGs had higher dN/dS ratios (8% showed dN/dS > 1) than the singletons (0.2% showed dN/dS > 1) and the genomic average, suggesting larger sequence divergence and weaker evolutionary constraints (Fig. 3B). In fact, we were able to identify positively selected amino acid sites among the protein sequences in a significant number of expanded OGs using the CodeML module of PAML (27) (Fig. 3C). Interestingly, in FBP, LEC, OAC, and zf-CCCH (containing CCCH-type zinc finger domain) genes, the sites never occurred in the identified domains, while other parts of the protein were subjected to positive selections (Fig. 3C and D). In GPCR_*Srz* family, many positively selected sites were mapped to the transmembrane helices and extracellular regions and might affect residues involved in ligand binding. A previous study analyzing a group of nine *Srz* genes in *C. elegans* reached similar conclusion (28).

Expression analysis showed that genes of the expanded and specific OGs are expressed in fewer tissue and at lower levels compared to singletons (Fig. S2), which is consistent with the findings for duplicate genes. Different gene families also showed enriched expression in different tissues. For example, NHR genes were highly expressed in seam cells, Caludin_3 genes (involved in forming tight junctions) in the intestine, and DUF19 genes in anal and body wall muscles (Fig. S2).

Because the *C. elegans* expanded and specific OGs contained many FBP, MATH, and LEC genes, which were known to be involved in stress response and innate immunity (29), we combined and analyzed the RNA-seq data for gene expression changes upon exposure to heat shock and five pathogens (see Methods for details). We found that genes in the *C. elegans* expanded and specific OGs were enriched (enrichment fold = 1.3) in the significantly upregulated genes but not the downregulated genes (enrichment fold = 0.7) (Fig. 3E and F). Strikingly, the average fold change (28.7) was much higher for the duplicate genes in the *C. elegans* expanded and specific OGs than the average fold change (2.1) of the singletons in the six stress conditions (Fig. 3G); the duplicate genes also appeared to respond to more than one conditions, whereas the singletons responded specifically to only one stress on average (Fig. 3H). These results support the findings in yeast that duplicate genes have higher plasticity and larger dynamic range in conditionally responsive expression (30). Nonetheless, in yeast, duplicate genes respond to specific stress and singletons respond more generally to many stress conditions (31); we found the opposite in *C. elegans*. In fact, using the WormExp gene set enrichment analysis (32), we found that the duplicate genes were also enriched in genes upregulated by the infection of many other pathogens (Fig. 3I), supporting the versatility of their functions in immune response.

### Functional diversification of the recently duplicated genes

Divergence in expression pattern is an indication of functional divergence. In fact, classical examples of subfunctionalization of duplicate genes often involve partitioned expression of the paralogs (2). Since most (81%) recently duplicated genes showed expression in at least one neuron, we examined their expression patterns in the nervous system using the single-cell transcriptomic data (22). From the *C. elegans* expanded and specific OGs, we first picked the OGs with only two *C. elegans* genes and found that the expressions of most paralogous pairs were highly divergent in the nervous system (Fig. 4A), indicating possible functional diversification. When examining OGs with more than two duplicate paralogs, we found significant divergence in expression patterns in general, despite some conserved expression in certain neurons (e.g., GPCR expression in ADL neuron and IRLD expression in the ASJ) (Fig. 4B). We reason that the genes in these rapidly expanding OGs may be generated by very recent duplications, so they still preserved some shared ancestral expression while acquiring new expression in other cells.

**Figure 4.**
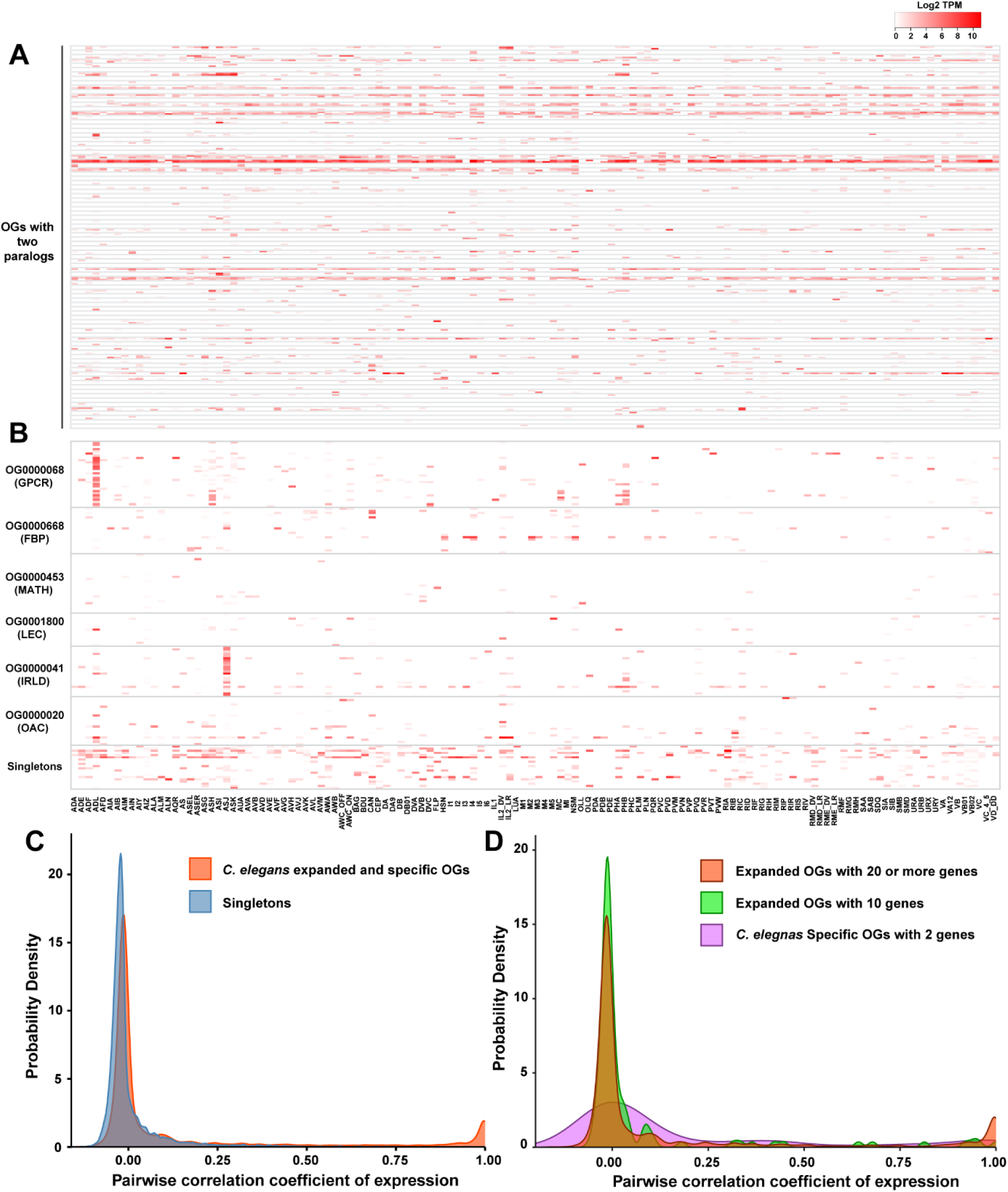
Expression divergence of recently duplicated genes. (A) The expression pattern of paralogous pairs in 88 *C. elegans* specific OGs that contained only two genes. OGs are separated by grey lines in the rows. The columns showed the expression levels in 128 neuron types of the nervous system. (B) The expression pattern of duplicate genes in representative *C. elegans* specific or expanded OGs with more than two genes. As a comparison, twenty singletons were randomly chosen from the singletons that are expressed in less than 50% of the neuron types. (C) The probability density distribution of Pearson correlation coefficient for pairwise comparison among paralogs in the same OGs. In total, correlation for 13,431 pairs were calculated for the C. elegans expanded and specific OGs; the same number of randomly chosen pairs of singletons were used as a control. (D) The probability density distribution of pairwise Pearson correlation coefficient of expression levels for paralogs in *C. elegans* specific OGs with two genes and *C. elegans* expanded OGs with 10 or at least 20 paralogs.

For more quantitative measurements, we calculated the pairwise correlation coefficients of the expression level for duplicate genes in the same OG and as a control for all singletons. From the probability density distribution, we found that most paralogs in *C. elegans* expanded and specific OGs had no correlation (the peak around 0) in expression, similar to the unrelated singleton pairs; some pairs showed high correlation (the peak close to 1) presumably because they were newly duplicated paralogous pairs that did not diverge yet (Fig. 4C). This idea is supported by the observation that rapidly expanding OGs that have many paralogs contained more pairs with strong correlation than the expanded OGs with fewer paralogs (Fig. 4D).

### Intraspecific structural variation of *C. elegans* genome

We next studied the fate of the recently duplicated genes during the intraspecific evolution of *C. elegans* genome using the sequence of 773 *C. elegans* wild strains (CeNDR 2020 release). We first called the structural variants (SVs) through a computational pipeline (Fig. 5A) and identified in total 74,360 high-quality SVs, including deletion, insertion, inversion, and duplication (Fig. S3A-C). We excluded >1 Mb variants since there are very few such variants and we cannot confirm their validity; similar exclusions were adopted by previous studies that called SVs from short-read sequencing data (33). In contrast, we were able to confirm seven randomly picked deletions ranging from 71 to 2392 bp by genotyping the wild strains that carry the deletion (Fig. S3D). Vast majority (92%) of the deletions were smaller than 2 kb.

**Figure 5.**
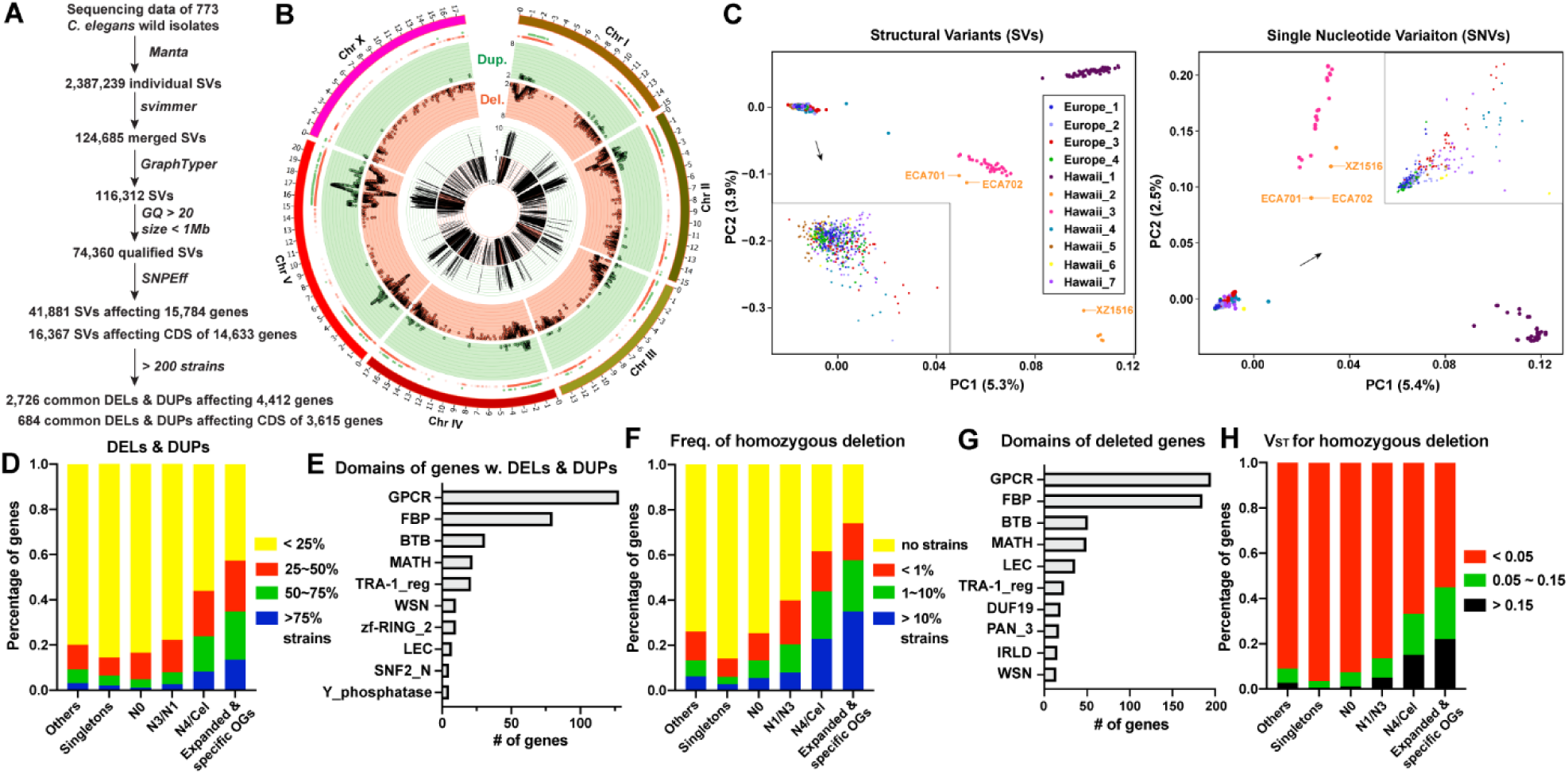
Structural variation among the 773 wild *C. elegans* strains. (A) A computational pipeline we used to calle the structural variants (SVs) among the *C. elegans* wild strains. (B) The distribution of deletions (DELs) and duplications (DUPs) across six chromosomes. The outmost ring showed the density of DELs and DUPs; the scatter plot in the middle ring showed the distribution of DELs and DUPs with each point representing the number (log2 transformed) within a 1,500-bp genomic window; the histogram in the inner ring showed the number of strains (log2 transformed) carrying those CNVs in the genomic window. Green represents duplication events and red for deletion events. (C) Principal component analysis for SVs and SNVs of 11 genetic groups. One point indicates one individual strain. Three highly divergent strains, XZ1516, ECA701 and ECA702 belonging the Hawaii_2 group were labeled. (D) The percentages of genes whose coding regions are affected by DELs or DUPs in different numbers of strains. (E) The top ten domains for the N4/Cel and *C. elegans* expanded &specific genes affected by DELs and DUPs in more than 75% of the strains. (F) The percentages of genes that carry homozygous deletions affecting their coding regions in different numbers of strains. (G) The top ten domains for the N4/Cel and expanded &specific genes carrying homozygous deletions in more than 10% of the strains. (H) The percentages of genes with V_ST_ values within the indicated ranges for the six gene groups. The V_ST_ value was computed by setting the copy number to 0 for homozygous deletions.

Among the SVs, the 61,003 deletions (DELs) and duplications (DUPs), are mostly located at the chromosomal arms, especially on chromosome V (Fig. 5B). Interestingly, majority of the recently duplicated genes were also found on the arms of chromosome V (Fig. 1C), implying that the same mechanism (e.g., non-allelic homologous recombination) that drove gene duplication may also lead to the generation of intraspecific DELs and DUPs.

Using the SV data, we analyzed the population structure of the 773 wild strains and identified 11 genetic groups (Fig. 4C and Table S7; see Methods for details). Principle component analysis showed the genetic divergence of three Hawaiian groups from the other eight groups. The SV-based population structure matched the SNV-based structure (Fig. 5C). Thus, our studies with the SVs support the findings of previous studies (34, 35) using the SNVs that some Hawaiian strains carry ancestral variants that are highly divergent, while other Hawaiian strains (Hawaii_4-7 groups) are admixed with the European groups.

### Recently duplicated genes are frequently deleted within the species

From the SV data, we found that duplications and deletions occurred at much higher frequencies in the recently duplicated genes and in the *C. elegans* expanded and specific OGs than in older duplicated genes and singletons, indicating more dynamic intraspecific evolution (Fig. 5D and E). More specifically, we found that 35% of the *C. elegans* expanded and specific OGs and 23% of the recently duplicated genes were homozygously deleted in more than 10% of the wild strains (Fig. 5F). As a comparison, only 3% of the singletons were deleted at the same frequency. Thus, the genes newly generated at the species level were often deleted among the individuals of the same species, suggesting the lack of evolutionary constrains and fast gene turnover. GPCR, FBP, BTB, MATH, LEC, and IRLD genes were among the top gene families that showed high frequency of homozygous deletion (Fig. 5G).

By calculating the fixation index (V_ST_) among the genetic groups, we found that homozygous deletions in recently duplicated genes showed much higher V_ST_ values than older genes and singletons, suggesting that the deletion of the young duplicate genes contribute to population structure (Fig. 5H). In fact, some of these genes were deleted in almost all strains of certain genetic groups (Fig. S4). These results suggest that the deletion of some duplicated genes might provide certain advantages for the population. One such example is a natural deletion in *srg-37*, which reduced pheromone sensitivity to increase population density (36). Interestingly, this *srg-37* (*DEL_94bp*) deletion occurred at much higher frequency in two genetic groups than in other groups (Fig. S5A).

### Young duplicate genes showed large intraspecific variations and signs of balancing and positive selection

By calculating population genetics statistics for both nonsynonymous SNVs and SVs (i.e., DELs &DUPs), we found that the recently duplicated genes and the genes in *C. elegans* expanded and specific OGs showed much larger polymorphisms than singletons and older duplicate genes (Fig. 6A-C and Table S8). We observed a clear trend for younger duplicate genes carrying larger intraspecific variations by comparing genes duplicated at N0, N1/N3, and N4/Cel stages.

**Figure 6.**
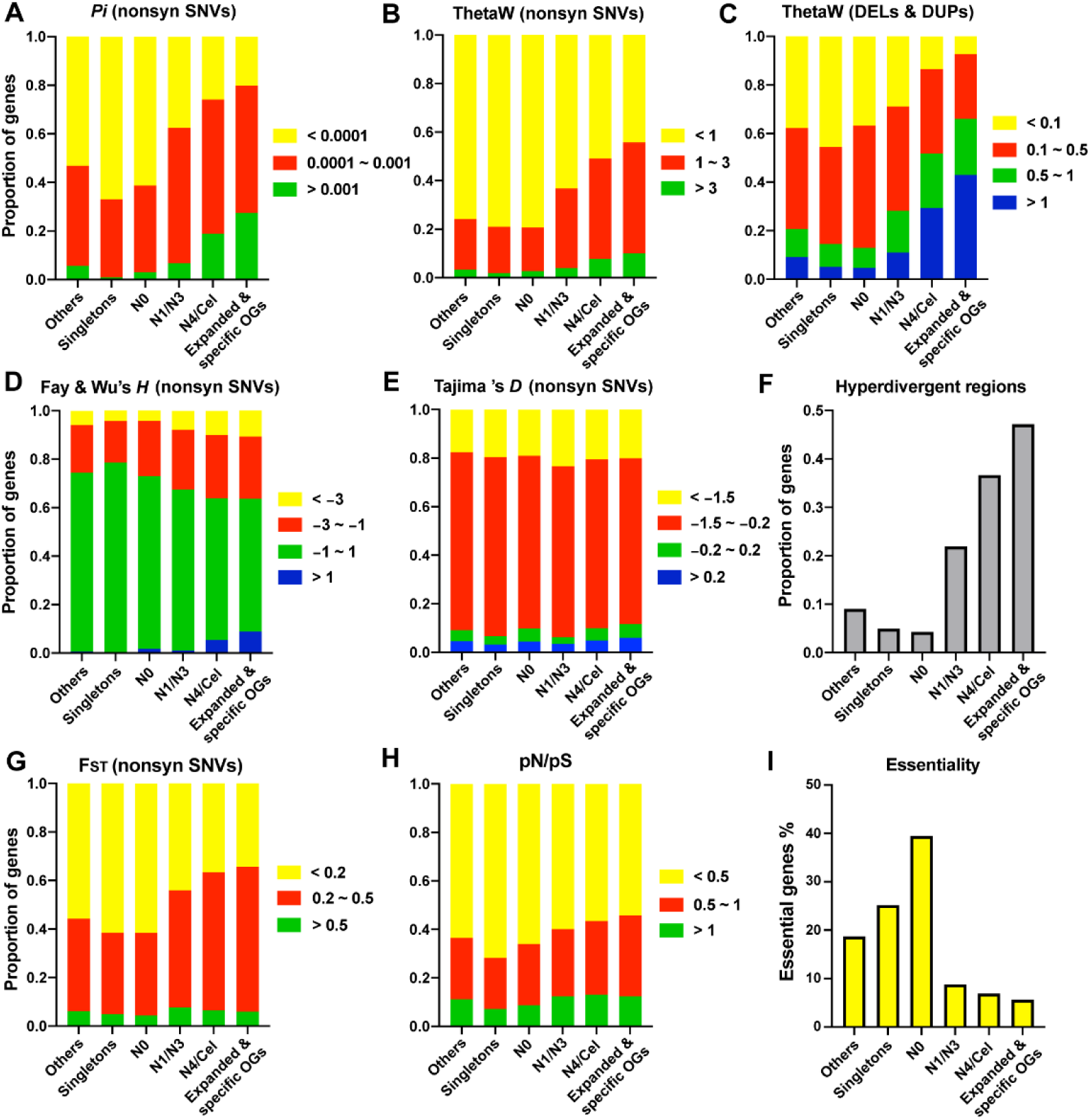
Intraspecific variations and molecular evolution of recently duplicated genes. (A-E) The percentages of genes with *Pi* (A), ThetaW (B-C), Fay &Wu’s *H* (D), and Tajima’s *D* (E) values within the indicated ranges for the six gene groups. Hawaii_2 strain XZ1516 was used as the outgroup for the calculation of *H*. (F) The percentage of genes located in the hyperdivergent regions as defined by Lee *et al*. (37). If a gene carries nonsynonymous variants in strains that have hyperdivergent genomic regions overlapping with the genomic location of the gene, we considered the gene to be in hyperdivergent regions. (G-H) The percentage of genes with F_ST_ (G) and pN/pS (H) values within the indicated ranges for the six gene groups. (I) The percentage of genes that were found to be essential genes based on previously published RNAi phenotypes.

The large genetic diversity in recently duplicate genes may result from not only weak purifying selection but also balancing selections, since we observe more genes among the young duplicate genes to show positive Fay and Wu’s *H* and Tajima’ *D* than singletons and older genes (Fig. 6D-E). Previous studies found that 20% of the *C. elegans* genome contains punctuated hyperdivergent regions that show large variations between the wild isolates and the reference strain and are maintained by balancing selection (37). Recently duplicated genes are highly enriched in these hyperdivergent regions (e.g., about 50% of the *C. elegans* expanded and specific genes fall into these regions) (Fig. 6F). The lack of evolutionary constrains and the balancing selection of certain alleles in the young genes may have allowed significant variations to accumulate in these regions and contributed to their divergence.

In addition to balancing selection, a significant percentage of the young duplicate genes showed signs of positive selection, indicated by the more negative Fay and Wu’s *H* and larger fixation index (F_ST_) for nonsynonymous SNVs compared to older genes and singletons (Fig. 6D and G). Accumulation of high frequency derived alleles in the recently duplicated genes and the fixation of these alleles in certain genetic groups may have contributed to adaptation. As examples, we observed the missense variant in some recently duplicated GPCR genes in high frequency only in specific groups [e.g., *srg-37* (H84Q) in the Hawaii_5 group, *sri-26* (L266F) in Hawaii_5 and Hawaii_6 groups, and *sri-24* duplication in Hawaii_1 and Hawaii_3 groups] (Fig. S5A-B). Conversely, the reference allele appeared to be prevalent in certain groups indicated by the very low frequency for the derived allele [e.g., *srg-34* (I26V) and *sri-26* (E70Q) in Europe_1 and Europe_4 groups]. Similar group-specific presence or absence of SNVs and SVs was also observed in other OGs that experience significant expansion in *C. elegans*, such as the FBP gene families (Fig. S5C). Interestingly, some of the missense variants had PROVEAN score below the -2.5 threshold [e.g., -3 for *srg-37* (H84Q) and -4.1 for *fbxa-224* (N32D)], suggesting that they may affect protein functions, and structural variants often inactivate the gene [e.g., *fbxa-222* (DEL_142bp)] or increase the copy of the gene [e.g., *sri-24* (DUP_9396bp)] (Fig. S5D). These results suggest that variations in the recently duplicated genes might significantly alter the biological function of certain pathways in specific genetic groups, leading to intraspecific adaptation.

Another way to assess positive selection is to calculate the ratio of nonsynonymous (pN) to synonymous (pS) polymorphism. Recently duplicated genes showed higher pN/pS ratios than genes duplicated at earlier times and singletons, suggesting that stronger positive selection and more relaxed purifying selection (Fig. 6H). Strong positive selection was confirmed by the observation that the interspecific dN/dS value was higher than the intraspecific pN/pS value for 26% of the genes in *C. elegans* expanded and specific OGs and 20% of the recently duplicated genes, compared to only 7% for singletons.

Relaxed purifying selection was confirmed by the lower percentages of essential genes in the recently duplicated genes (6.9%) and the *C. elegans* expanded and specific OGs (5.6%) compared to genes duplicated earlier (39% at N0) and singletons (25%) (Fig. 6I). This result helps resolve a controversy for the essentiality of newly arisen genes. Chen *et al*., found that 30% of young genes (originated within the last 35 million years) in *Drosophila* were essential for viability based on RNAi results (7), whereas Kondo *et al*., in a later study examined knockout mutants of the young genes and found that most of them were not lethal (38). Our findings suggest that recently duplicated genes may be maintained through balancing and positive selection but rarely develop essential functions. Interestingly, genes duplicated at ancient times (N0; >130 mys ago) appeared to have higher percentage of essential genes than the singletons (Fig. 6I), indicating that duplicate genes could indeed develop strong essentiality over a long time.

### Copy number analysis support the dynamic intraspecific evolution of young duplicate genes

Finally, we studied the intraspecific evolutionary dynamics of duplicate genes using read depth-based copy number (CN) analysis. We applied CNVpytor, one of the most updated CN calling algorithms (39), to the whole-genome sequencing data of the wild isolates (median coverage is 36X) and identified 104 deletions and duplications ranging from 5 to 120 kb and affecting 658 protein-coding genes (Fig. 7A and Table S9). Most of these copy number variations are located at the chromosomal arms (Fig. 7B). Using the variance of CNs among the 773 wild strains to measure the CN diversity, we found that the recently duplicated genes showed larger CN variability than singletons and older duplicate genes (Fig. 7C). The genes that showed the largest variance in CNs are mostly FBP, GPCR, BTB, NHR genes, etc. (Fig. 7D).

**Figure 7.**
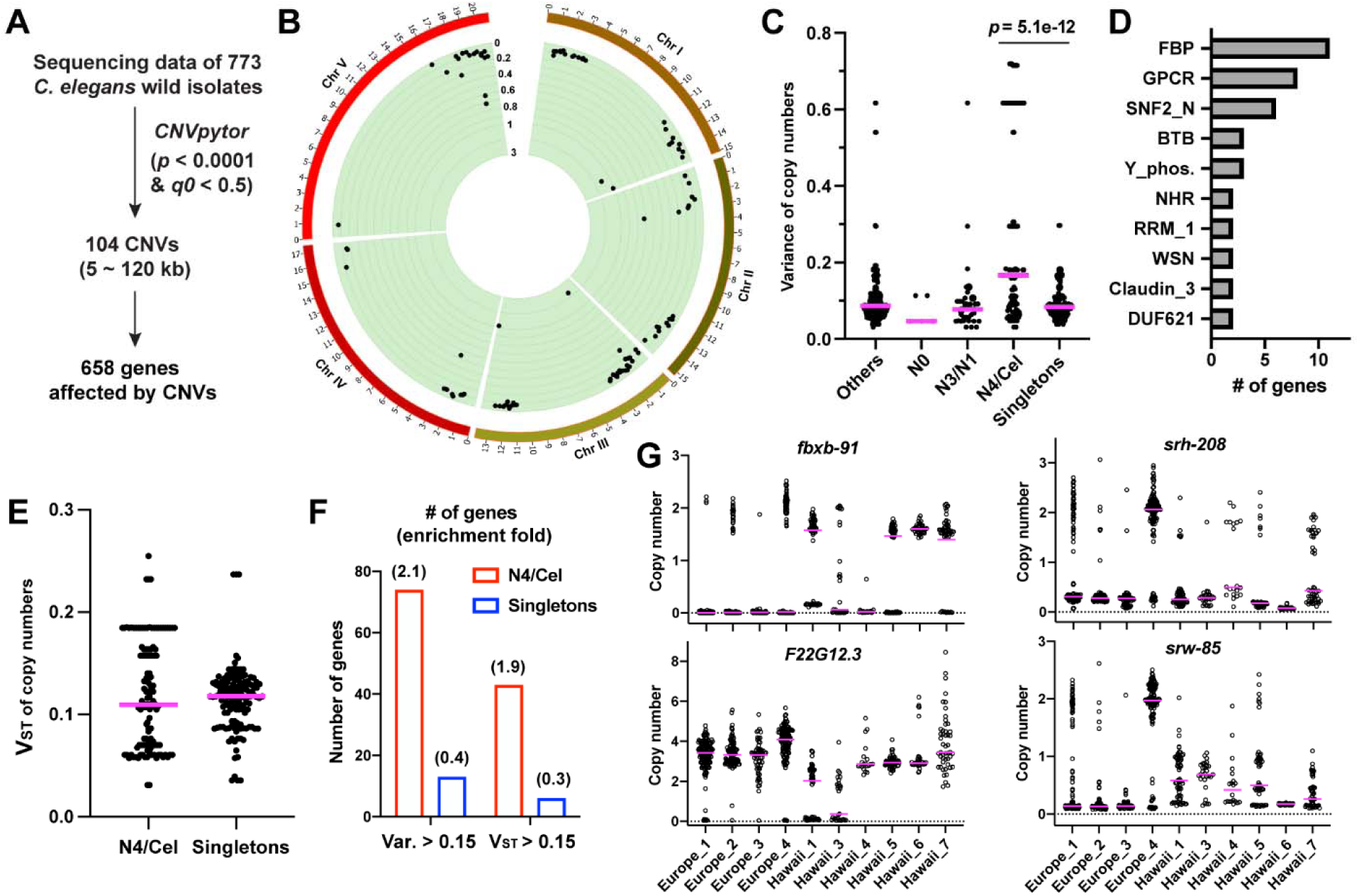
The copy number analysis of the wild isolates. (A) The pipeline we used to call the copy number variations (CNVs) from the whole genome sequencing data of the wild strains. (B) The circos plot showing the distribution of the 104 CNVs called by CNVpytor and the copy number variance (Var) among the wild strains. Chromosome X has no detectable CNVs and is not shown. (C) The Scatter plot showing the variance of copy numbers for genes carrying CNVs in each group. (D) The top ten domains for the 125 recently duplicated genes (N4/Cel genes) affected by CNVs. (E) The Vst distribution for 125 N4/Cel genes and 135 Singletons affected by CNVs. (F) The enrichment fold for N4/Cel genes and singletons in the genes with Var > 0.15 or V_ST_ > 0.15. (G) Representative FBP and GPCR genes with high V_ST_ and Var values. One circle indicates one strain in the genetic group and the purple line indicates the median copy number in the group.

Although the V_ST_ statistics for the CN differentiation among the genetic groups overall did not show significant difference between genes duplicated at N4/Cel and singletons, the N4/Cel genes are more enriched than singletons among the genes with relatively higher (> 0.15) V_ST_ values (Fig. 7E and F). This result suggests that at least some young duplicate genes tend to show more CN difference among the genetic groups than within the groups. For example, the FBP gene *fbxb-91* has 0 copies in most strains in Europe_1, Europ_2, and Hawaii_4 groups, but have normal copy numbers in 100% of the Hawaii_6 strains (Fig. 7G). Similar observations were made for another FBP gene *F22G12*.*3*, and two GPCR genes, *srh-208* and *srw-85*. CN difference among the groups might indicate positive selection by gene dosage effects, while large CN variance could also result from balancing selection. In general, the analysis of the CN variations supports the results with SNVs and SVs and confirms the dynamic intraspecific evolution of recently duplicated genes among the *C. elegans* wild isolates.

## Discussion

Although several previous studies investigated the adaptive evolution of duplicate genes at the species level through sequence comparison of homologs and paralogs (6, 40, 41), very few studies examined the evolutionary force that shaped the fate of duplicate genes at the individual level within a species. To bridge this gap, we analyzed both interspecific sequence divergence and intraspecific variation of the recently duplicated genes in *C. elegans* and found signs of positive selection acting at both the macroevolution and microevolution scales. Positively selected sites were found among the paralogs derived from recent duplication, which might drive functional divergence; meanwhile, the accumulation and positive selection of variants in the recently duplicated genes significantly contributed to the population structure, indicating a role in local adaptation. The Walsh model suggests that a newly duplicated gene can quickly evolve a novel and important function by accumulating advantageous mutations (42), but new function of the duplicate genes can only be acquired in large populations if the mutations are rare. Since *C. elegans* has a small effective population size (43, 44), our empirical evidence for unexpectedly prevalent positive selection acting on the recently duplicated genes suggest that positive selection of advantageous mutations may be an underestimated force driving the conservation of duplicate genes. In fact, this idea is supported by work in Drosophila and mammals, where a large percentage of duplicate genes are retained by positive selection and neofunctionalization (5, 6).

By comparing the young and old duplicate genes for expression pattern, selective pressure, and essentiality, our work led to a hypothesis for the fate of a duplicate gene in both short and long term (in evolutionary time scales). Shortly after the gene birth, relaxed purifying constraint allows the duplicate genes to accumulate mutations. High recombination rate at the chromosomal arms help maintain large genetic diversity in these genes and enable them to explore the evolutionary space until they acquire adaptive variants that are subjected to balancing or positive selection. This process occurs at both the population and the species levels. At this stage, the young genes are rarely essential for organismal development and are expressed in a few specific tissues at low levels. Following the preservation of young duplicate genes through positive selection, it takes at least tens of millions of years for the genes to broaden expression patterns, evolve important functions in vital pathways, and finally be retained by purifying selections. In fact, duplicate genes become as essential as the single-copy genes in the long term, suggesting that the accumulation of novel function may eventually convert duplicate genes into functional singletons.

What are the functions of the young duplicate genes before it evolves essential functions? In *C. elegans*, we found that genes involved in chemosensory perception (e.g., GPCRs), protein degradation pathways (e.g., FBP, MATH, and BTB), and pathogen detection (e.g., LEC) are specifically enriched in the recently duplicated genes. Interestingly, GPCRs, FBP, MATH, and LEC genes all play important roles in innate immune response (29, 45). Indeed, we found that although the basal expression of the recently duplicated genes is low, their expression is highly responsive for pathogenic infection with large dynamic range; and they respond to multiple stress conditions. Thus, although the recently duplicated genes do not participate in development and housekeeping functions, their conditional expression upon external perturbation, such as heat stress and infection, may contribute to the adaptability of the organism in a changing environment. Being consistent with this idea is the significant expansion of the GPCR family, which are the chemoreceptors that detect not only external odors and chemicals but also internal peptides (46, 47). A recent study hypothesized that the nuclear hormone receptors (NHRs) may control the expression of GPCRs for the integration of sensory information in sensory neurons (48). Interestingly, we found that the NHR families were also significantly expanded in *C. elegans*. Thus, the co-expansion of GPCRs and NHRs families may facilitate the establishment of an NHR-GPCR regulatory network to drive adaptation behaviors. Overall, our study provides important insights into the function of the young duplicate genes in sensing and responding to external signals and perturbations.

## Methods

### Construction of orthogroups (OGs) and the identification of duplicate genes

From the genome assemblies of twenty-one *Caenorhabditis* species available in the WormBase ParaSite database (Table S1), we selected eleven species based on the assembly contiguity (N50 > 200,000) and the completeness of genome assembly and annotation (BUSCO assembly > 90%). We did not include *C. brenneri* due to the reported high heterozygosity (49). The longest protein isoform encoded by each gene in the eleven species was extracted and used for constructing OGs using the OrthoFinder v2.5.4 (19) with default parameters (Table S2). The species tree was inferred from all genes using the STAG algorithm (50) and then rooted using the STRIDE algorithm (51) for identifying gene duplication events.

Previously, Cutter et al., measured the divergence times by estimating neutral mutation rate among a few *Caenorhabditis* species (52). To estimate the divergence time on the phylogenetic tree of the eleven *Caenorhabditis* species, we applied the “r8s” program (53) on the tree and set the median divergence time between *C. elegans* and *C. briggsae* as 60 million years ago (according to TimeTree (54)) for calibration. Duplication events at each branch in the species tree were identified using the Duplication-Loss Coalescence algorithm (20) implemented in OrthoFinder; duplication events with support possibility lower than 0.5 were discarded. Genes with duplication events in more than one branch were assigned to the most recent one. Genes with duplication events within 60 million years were considered as recently duplicated genes.

### Identification of *C. elegans* expanded and specific OGs

To identify significantly expanded and contracted OGs, we used the gene birth-death model implemented in CAFE v4.2.1 (26) (https://hahnlab.github.io/CAFE/manual.html) to analyze the evolution of gene family size on 24,422 OGs. Species tree constructed by OrthoFinder was used as the input for CAFE. A single birth and death parameter λ was estimated based on the estimated divergence time (60 million years ago) between *C. briggsae* and *C. elegans* by TimeTree (http://timetree.org/) (54). The rapidly evolving gene families at each branch on the tree were then identified as significantly expanded and contracted OGs and were used for further analysis. *C. elegans* specific genes refer to the genes within the OGs that only contained *C. elegans* genes. In total, we obtained 1,128 genes from 71 *C. elegans* expanded OGs and 1,123 genes from 264 *C. elegans* specific OGs. Genes in these OGs were subjected to gene ontology analysis using the GO tool and the enrichment of functional modules was visualized using ClueGO App in Cytoscape (55).

### Structural variant (SV) calling, copy number (CN) analysis, and calculation of population genetics statistics for *C. elegans* wild strains

We used the quality-filtered and aligned whole genome sequencing data for the 773 *C. elegans* wild strains from the *Caenorhabditis elegans* Natural Diversity Resource (CeNDR) website (10) (https://www.elegansvariation.org/) and applied Manta v1.6.0 (56) to call SVs on the alignment libraries of the wild strains with WBcel235 as the reference genome). We merged called genotypes for all samples using svimmer (https://github.com/DecodeGenetics/svimmer) and applied GraphTyper 2.0 (57) to recall and genotype SVs. We then removed the SV larger than 1 Mb and set the genotypes with Genotype Quality (GQ) smaller than 20 or with “FAIL” tag as missing genotype. The genotyped VCF file was then annotated using SNPEff v4.3t (58) with the Ensembl WBcel235 genome assembly as the reference. To characterize the copy number variation among *C. elegans* wild isolates, we applied CNVpytor, which is a read depth (RD)-based algorithm and an updated version of the widely used CNVnator (39, 59). We assigned the copy numbers of the genes affected by duplication and deletion events identified by CNVpytor.

To calculate population genetics statistics, we used PopGenome (60) to calculate *Pi* (61), Watterson’s theta (ThetaW) (62), Hudson’s F_ST_ (63), Tajima’s *D* (64), and Fay and Wu’s *H* (65) for nonsynonymous SNVs of each gene. A highly divergent strain, XZ1516, was used as the outgroup for calculating *H*. We calculated ThetaW and V_ST_ (66) for DUPs and DELs using similar formulas as for SNVs.

### Other methods

Detailed information about the above methods and other methods (including protein domain prediction, collinearity analysis, phylogenetic tree construction, identification of positively selected sites, RNA-seq analysis for stress response, expression pattern analysis, population structure analysis, population genetics statistics calculation, gene essentiality analysis, etc.) can be found in *SI Appendix*.

## Acknowledgement

We thank the WormBase Parasite for sharing genomic data of *Caenorhabditis* species, the *Caenorhabditis elegans* Natural Diversity Resource (CeNDR) at Northwestern University for sharing genomic data of the *C. elegans* wild isolates, and the *C. elegans* Neuronal Gene Expression Map &Network (CeNGEN) for sharing single-cell transcriptomic data. This study was supported by funds from the Research Grant Council of Hong Kong [ECS 27104219, CRF C7026-20G, and GRF 17107021], the Food and Health Bureau of Hong Kong [HMRF 07183186], the National Natural Science Foundation of China (Excellent Young Scientists Fund for Hong Kong and Macau), and the seed fund from the University of Hong Kong [201910159087 and 202011159053] to C.Z. Computational works were performed using research computing facilities offered by Information Technology Services at the University of Hong Kong.

## Notes

### Competing Interest Statement

The authors have declared no competing interest.

