## Supplemental text for "Dynamic evolution of recently duplicated genes in *Caenorhabditis elegans*"

**This PDF file includes:**

Supplementary text (SI Methods)
Tables S1-S10 legends
Figures S1-S6

### SI Methods

#### Protein domain prediction and PROVEAN score

We used PfamScan to search for the potential domains for each gene in Pfam-A protein database (1). HMM file for specific domain was downloaded from the Pfam website (https://pfam.xfam.org/) and was used to search against the protein sequence files of the eleven species using hmmsearch. TMHMM server was employed (https://services.healthtech.dtu.dk/service.php?TMHMM-2.0, last accessed March 1, 2022) to predict the transmembrane domain (TM) and the intracellular and extracellular regions of GPCR proteins. One gene often carries more than one domain. Thus, we merged genes carrying the F-box, FTH, FBA_2 and HTH_48 domains as F-box proteins (FBPs) and merged all GPCR subfamilies as GPCRs (see Table S5).

The potential functional alteration of nonsynonymous mutations was predicted using the PROVEAN (Protein Variation Effect Analyzer) web server (2) (http://provean.jcvi.org/seq_submit.php). A PROVEAN score smaller than -2.5 was considered to potentially change protein functions.

#### Collinearity analysis

We used MCScanX (3) to reveal the history of GPCR, FBP and NHR gene family expansions and analyze the duplication types by identifying putative homologous chromosomal regions. E value threshold of 1e-5 was used to construct collinearity background. Genes in the recently duplicated GPCR, FBP and NHR families were used as the input homologs for the collinearity analysis.

#### Phylogenetic tree construction

Protein sequences from the same OG or a list of proteins containing the same domain were aligned using MUSCLE v3.8.31 (4). The aligned sequences were trimmed by clipkit (5). We then used iqtree v2.1.3 (6) to construct the phylogenetic tree by maximum likelihood with 1000 bootstraps. The parameter –m MFP was used to determine the best-fit substitution model. The ultimate tree file was modified using iTOL (7).

**Identification of positively selected sites and calculation of dN/dS (ω) ratios**

Protein sequences from the same OG were first aligned by MUSCLE and then aligned by codon alignment program in MegaX (8). Gaps were removed after the alignment. We then ran the M0, M7 and M8 models in the CodeML program of PAML v4.9j (9) to identify positively selected sites among paralogs. We tested the positive selection by comparing twice the log-likelihood difference between M7 and M8 models with a $\chi^2$-distribution in the likelihood ratio test (LRT). The dN/dS values for each gene were computed using a free-ratio model which allows ω to vary along branches (NSsites = 0, model = 1).

**RNA-seq analysis for stress response**

RNA-seq data for the six stress conditions were downloaded from multiple previous studies (we chose studies with at least two replicates for accuracy). The stress conditions include heat shock (10), and infections by *Pseudomonas aeruginosa* (PA14) (11), *Bacillus thuringiensis* (12), *Serratia marcescens* (13), *Nematocida parisii*, and Orsay virus (14). Reads were aligned to the WBcel235 reference genome of *C. elegans* by STAR v2.7 (15). Gene count was computed using STAR option --quantMode GeneCounts for RNA-seq data under different stress treatments. The significantly differentially expressed genes under these stresses were identified using DESeq2 (16). To identify more pathogens that may induce expression of the recently duplicated genes, we analyzed the genes in *C. elegans* expanded and specific OGs using the WormExp under the "Microbes" category (17).

**Expression pattern analysis in *C. elegans***

We obtained the single cell transcriptomic data for most cell types of *C. elegans* from the *C. elegans* Neuronal Gene Expression Map & Network (CeNGEN) (18) by downloading the "all cells unfiltered" gene expression data for eleven tissues and 128 individual neuron types. Expression level was measured as log10 of transcripts per million (TPM). For the nervous system expression, we took the mean value of the 128 neuron types as the representative expression value for each gene.

The heat-map was plotted using heatmap.2 function in R. The expression was clustered by Dendrogram.

**Structural variant (SV) calling for *C. elegans* wild isolates**

To analyze the genetic diversity among the *C. elegans* wild isolates, we downloaded the quality-filtered and aligned whole genome sequencing data for the 773 *C. elegans* wild strains from the *Caenorhabditis elegans* Natural Diversity Resource (CeNDR) website (19) (https://www.elegansvariation.org/). We then applied Manta v1.6.0 (20), an assembly-based algorithm, to call SVs on the alignment libraries of the wild strains. The reference Genome assembly (WBcel235) of *C. elegans* was downloaded from Ensembl Genome Browser. Each inversion event was reported as multiple breakends in Manta, so we used a script provided by Manta (convertInversion.py) to convert these breakends into single inversion events. When discovering SVs, Manta may underestimate the joint calling frequencies. To resolve this, we subsequently merged called genotypes for all samples using svimmer (https://github.com/DecodeGenetics/svimmer) and applied GraphTyper 2.0 (21), which builds pangenome reference to genotype SVs. This algorithm re-aligned short reads to a pangenome reference and gives three types of genotyping format: coverage, breakpoint, and aggregated model. We extracted and used the aggregated model for further analysis. We then removed the SV larger than 1 Mb and set the genotypes with Genotype Quality (GQ) smaller than 20 or with "FAIL" tag as missing genotype. After the quality control step, the ECA551 has no qualified genotype for any SV (likely due to the low quality of the alignment file for the strain), so the final SV dataset contained only 772 strains. The genotyped VCF file was then annotated using SNPEff v4.3t (22) with the Ensembl WBcel235 genome assembly as the reference. We used Circos (23) to plot the distribution of deletions and duplications and the number of strains carrying these SVs.

The size distribution of the four types of SVs were shown in Fig. S3A-C. 85% of the SVs we identified were deletions, and 92% of the deletions were smaller than 2 kb. Thus, to validate the SVs called, we randomly pick seven deletions ranging from 71 to 2392 bp and genotyped the wild strains carrying these deletions through worm PCR and Sanger sequencing of the PCR product (Fig. S3D). Genotyping primers are listed in Table S10. Deletions and insertions in the SV dataset were bigger than 50 bp (< 50 bp deletions and insertions) were included in the SNV dataset as small indels). In our analysis, we also confirmed previously identified natural SVs. For instance, previous studies found a 94-bp deletion in *srg-37* (24) and a 159-bp deletion in *drh-1* among the wild strains (25) and we confirmed these two deletions in our SV calling. Moreover, we compared the SV outputs of both Manta+GraphTyper and CNVpytor (see below) and found that 103 out of the 104 SVs called by CNVpytor are overlapped with these identified by Manta+GraphTyper.

**Population structure among the wild isolates**

We applied Admixture (1.3.0) (26) on the genotyped VCF file for structural variants (SVs; called in this study) and single nucleotide variants (SNVs; CeNDR 2020 release downloaded from https://www.elegansvariation.org/data/release/20200815) to analyze the ancestral proportion of wild *C. elegans* strains (Table S7). Prior to that, we used PLINK (v1.9) (27) to prune the VCF files to remove variants with high pairwise linkage disequilibrium (LD) (--indep-pairwise 50 1 0.9). We set the number of genetic groups to be 11 because the cross-validation (CV) error reached a low level when K = 11 and remained at the similarly low level when K > 11 (Fig. S6). Moreover, we compared the strains distribution in particular genetic groups for SV- and SNV-based structures and found that on average over 70% are overlapped between them, suggesting that the population structures were similar for SVs and SNVs. Principle component analysis (PCA) on the VCF files for both SVs and SNVs were performed using the R package “SNPRelate” (28). The VCF file were converted to gds format using the function *snpgdsVCF2GDS* in the “SNPRelate” package for the analysis.

**Calculation of population genetics statistics for *C. elegans* wild strains**

For SNVs, we used PopGenome (29) to calculate *Pi* (30), Watterson's theta (ThetaW) (31), Hudson's $F_{ST}$ (32), Tajima's *D* (33), and Fay and Wu's *H* (34) for each gene. A highly divergent strain, XZ1516, from the Hawaii_2 group was used as the outgroup for the calculation of *H* (the other six Hawaii_2 strains were excluded when calculating *H*). We computed the ratio (pN/pS) of nonsynonymous over synonymous difference based on previous formula (35). ThetaW and $V_{ST}$ (36) for DUPs and DELs are computed as the formulas below.

$$\theta_w = \frac{K}{\sum_{i=1}^{n-1} \frac{1}{i}}$$

Where K is the number of segregating sites and n is the number of chromosomes.

For the calculation of $F_{ST}$ and $V_{ST}$, we removed 68 strains whose highest ancestral proportion were less than 0.5 and the Hawaii_2 group, which contained seven highly divergent strains. The remaining 698 strains were divided into 10 groups (Table S7). Pairwise $F_{ST}$ and $V_{ST}$ values were computed based on the grouping.

For example, the formula to calculate $V_{ST}$ is:

$$V_{ST} = \frac{V_{total} - (V_{pop1} \times N_{pop1} + V_{pop2} \times N_{pop2})/N_{total}}{V_{total}}$$

Where $V_{total}$ is the total variance, $N_{total}$ is the total population size among all populations. $V_{popx}$ and $N_{popx}$ are the variance and population size for each population, respectively.

**Gene essentiality analysis**

Gene essentiality analysis was performed based on the previous RNAi phenotype data for lethality (37). Genes whose RNAi knockdown led to any of the "lethal", "embryonic_lethal", "adult_lethal", "embryonic_terminal_arrest_variable_emb", "embryonic_lethal_late_emb, larval_lethal", "larval_arrest", "late_larval_lethal", "late_larval_arrest", and "one_cell_arrest_early_emb" phenotypes on WormBase WS277 were deemed to be essential. Essential genes in singletons, *C. elegans* expanded & specific OGs, and duplicate genes at N0, N1/N3 and N4/Cel branches were counted, and percentages were calculated.

### Copy number (CN) analysis

To characterize the copy number variation among *C. elegans* wild isolates, we applied CNVpytor, which is a read depth (RD)-based algorithm and an updated version of the wildly used CNVnator (38, 39). We also included the B-allele frequency likelihood information from the SNV data, which is complementary to RD signal, for calling the copy number variation through CNVpytor. The median coverage depth for the 773 strains from CeNDR is about 36. The bin size was set as 1,000 as CNVpytor suggests. We made a joint call set by merging calls from different samples with > 50% reciprocal overlaps. We then filtered the calls to keep the ones with $p < 0.0001$ and $q0 < 0.5$. The resultant file was annotated to identify the genes affected by the variations, and the copy numbers of the affected genes were assigned based on the duplication or deletion events affecting the region.

**Table S1. Summary of the genome assembly quality for twenty-one *Caenorhabditis* species.** Genomic data were downloaded from WormBase ParaSite website (https://parasite.wormbase.org/ftp.html). Contig number was counted based on the genome assembly file. The genome assemblies of eleven species were qualified for follow-up analysis (highlighted in yellow).

**Table S2. Orthogroups (OGs) assignment for genes in the eleven *Caenorhabditis* species.** The longest transcript for each gene in the eleven species was used to construct OGs. In total, 24,422 OGs were constructed.

**Table S3. Domain of duplicate genes in eleven species.** Domain information for the genes duplicated in different time periods were obtained by searching against the Pfam database using hmmsearch.

**Table S4. Duplicate genes in *C. elegans* genome.** *C. elegans* genes generated by duplication events at different branches of the phylogenetic tree (including N0, N1/N3, and N4/Cel branches). *C. elegans* singletons were also included for comparison. Calculated *dN/dS* values for each gene can be found in the table.

**Table S5. The number of significantly expanded and contracted OGs with specific domains in the eleven *Caenorhabditis* species.** OGs were grouped together based on the molecular or functional similarities of the domains identified in the OGs. Some OGs contained multiple domains, so the number of OGs in specific gene families may be smaller than the sum of the OGs with different domains.

**Table S6. Genes in *C. elegans* expanded and specific OGs with domain information.**

**Table S7. Admixture proportion of SNVs and SVs among *C. elegans* wild strains.** The table contains two sheets for the ancestral proportion of each strain calculated using SNV and SV datasets at K = 11, respectively. Grouping of the strains were done according to the population structure based on SNV dataset, which contains much more variants than the SV dataset.

**Table S8. Population genetics statistics for different types of genes.** Various population genetics statistics were calculated for every gene in *C. elegans* genome. Genes were marked for the groups they belong to. Duplicate entries were made for genes belong to both N4/Cel group and the *C. elegans* expanded and specific genes. *Pi*, thetaW, Tajima's *D*, Fay and Wu's *H*, $F_{ST}$, and pN/pS were calculated using the SNVs, while thetaW_DELs&DUPs was computed using deletions and duplications in the SV dataset (see methods for details).

**Table S9. Copy number variation of 658 genes among the *C. elegans* wild isolates.** Copy numbers of 658 genes affected by the 104 copy number variations called by

CNVpytor were listed for the 773 *C. elegans* wild strains. $F_{ST}$ values were calculated for each of the 658 genes.

**Table S10. Primers for validating seven randomly picked deletions.**

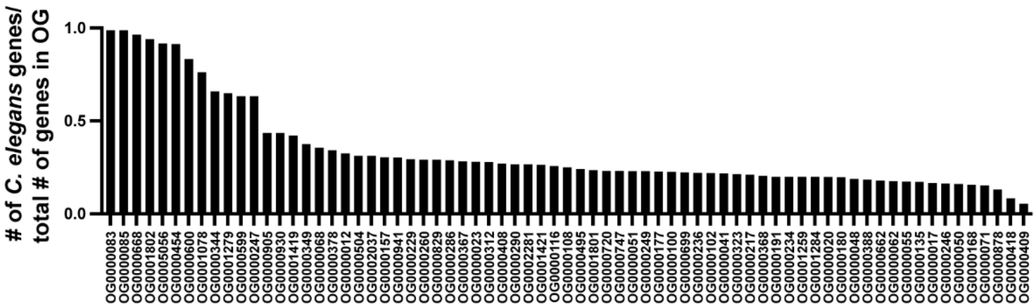


**Figure S1. The percentage of *C. elegans* genes in the 71 *C. elegans* expanded OGs.**

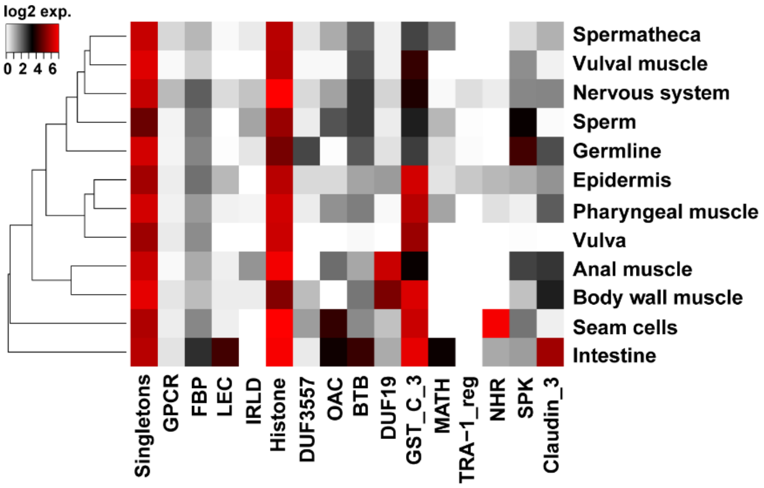


**Figure S2. Tissue expression patterns of the *C. elegans* expanded and specific genes with the top ten domains.** Expression data are presented as log2 transformed TPM (transcripts per million) values obtained from CeNGEN. For the nervous system, the mean value of the expression in 128 individual neuron types was used. Expression values were clustered by Dendrogram.

**A**

| | Median size (bp) | # of SV | |
|---|---|---|---|
| All SV | 149 | 74360 | |
| Deletion | 152 | 53835 | |
| Duplication | 1510 | 7168 | |
| Insertion | 310 | 8390 | |
| Inversion | 316 | 3490 | |
| | Median size (bp) | # of SV | # of genes affected |
| SV affecting genes | 187 | 41881 | 15784 |
| Deletion | 161 | 31149 | 13563 |
| Duplication | 2513 | 5562 | 9933 |
| Insertion | 95 | 2466 | 1931 |
| Inversion | 440 | 2252 | 5684 |
| | Median size (bp) | # of SV | # of genes affected |
| SV affecting CDS | 1317 | 16367 | 14633 |
| Deletion | 618 | 10357 | 11660 |
| Duplication | 5416 | 3793 | 9690 |
| Insertion | 129 | 736 | 694 |
| Inversion | 1351 | 1410 | 5205 |

**C**

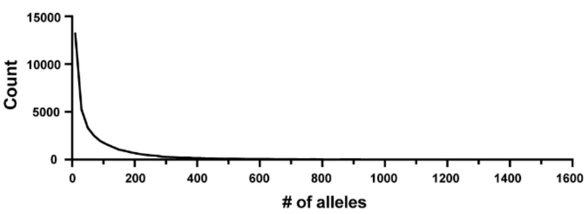


**D**

| Gene | Deletion Position | Predicted Deletion Size | Confirmed Deletion Size |
|---|---|---|---|
| *srp-6* | V: 7383315 | 2392 | 2392 |
| *zim-2* | IV: 10555639 | 2124 | 2124 |
| *pals-25* | III: 134629 | 1442 | 1538 (del + ins) |
| *lin-11* | I: 10253327 | 153 | 152 |
| *vab-3* | X: 10506065 | 131 | 131 |
| *ast-1* | II: 4134000 | 77 | 77 |
| *nhr-67* | IV: 11171755 | 71 | 65 |

**B**

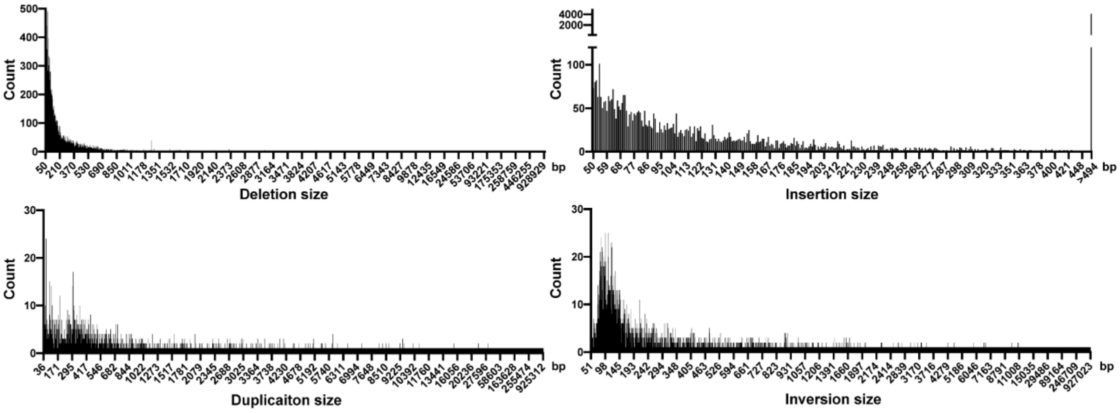


**Figure S3. A summary of SVs identified from 773 *C. elegans* wild strains.** (A) The number and median size of the four types of SVs, including deletion, duplication, insertion, and inversion, and the number of genes affected by these SVs. (B) Size distribution of deletions, insertions, duplications, and inversions. For insertions bigger than 494 bp, the exact size could not be obtained from the results of SV calling. Thus, they were binned together as > 494 bp. Deletions and insertions smaller than 50 bp were classified as small indels in the SNV dataset and thus excluded from the SV dataset. (C) The distribution of the allele frequency of the deletions (DELs) and duplications (DUPs) that affect the coding regions of genes among the 773 wild strains (1546 alleles in total). Most of the DELs and DUPs have very low allele frequencies. (D) Experimental validation of seven deletions ranging from 71 to 2392 bp in size by genotyping and Sanger sequencing.

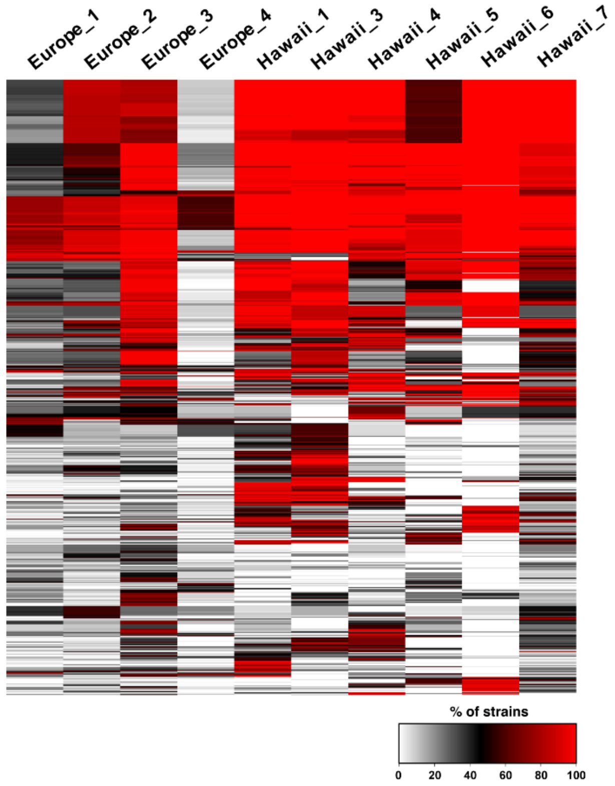


**Figure S4. Frequency of homozygous deletion in the recently duplicated genes in the ten genetic groups.** N4/Cel genes and genes in the *C. elegans* expanded and specific OGs with homozygous deletions in more than 10% of the 773 strains were plotted, and each row indicates one gene. Hawaii_2 group containing only seven strains was not included in the analysis.

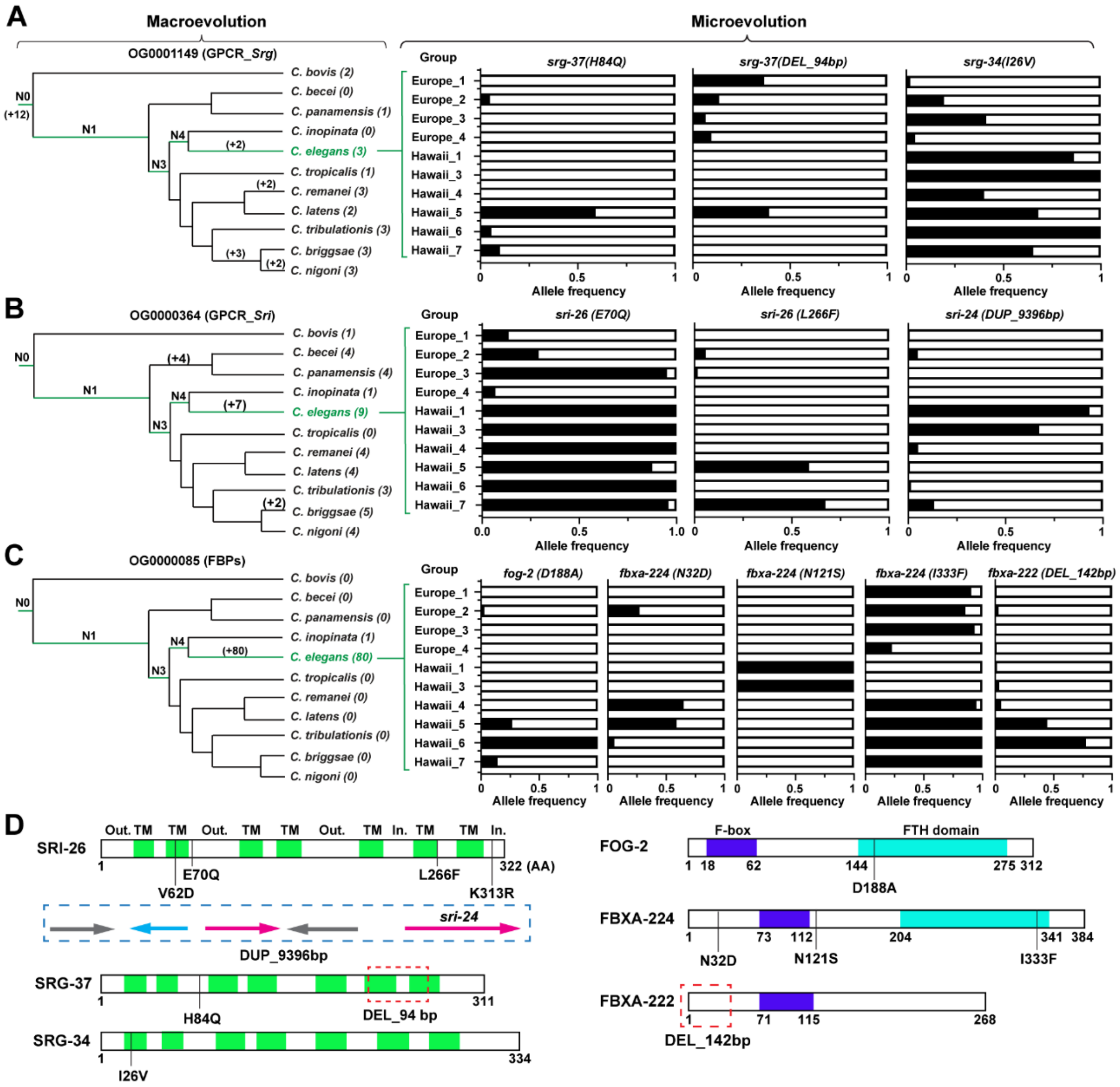


**Figure S5. Variations in recently duplicated genes tend to be fixed in specific genetic groups.** (A-C) The number (in parentheses) of genes generated by duplication events at different branches of the phylogenetic tree was shown for representative GPCR and FBP OGs. The allele frequencies of variations in the genes of these OGs in the ten genetic groups were shown in the right panels. (D) High frequency alleles were mapped to the domain structure of three presentative GPCRs and three FBPs. Green, the predicted transmembrane domain; blue, F-box domain; cyan, FTH domain. Duplication (blue dashed rectangle) and deletion (red dashed rectangle) were also indicated on the gene structure.

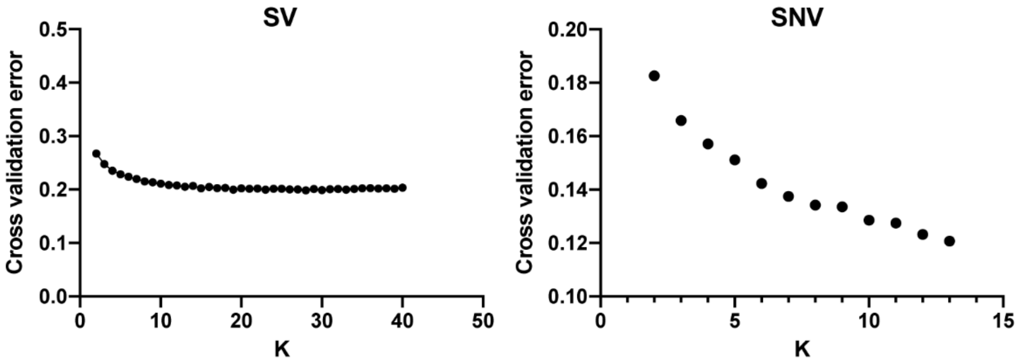


**Figure S6. The cross-validation (CV) error for population structure analysis.** CV errors for the number of population (K) ranging from 1 to 40 for structural variants (SVs) and from 1 to 13 for single nucleotide variants (SNVs). The CV errors were obtained from Admixture results of the population structure. Variants with high pairwise linkage disequilibrium were removed before the analysis.
